# Global Proteome Remodelling in *Rhodococcus jialingiae* RS1 to Decipher its Plant Growth-Promoting and Biofertilizer Properties: Gene Identification for Transgenics

**DOI:** 10.64898/2026.05.11.724437

**Authors:** Shakeel Ahmed Mohammed, Adesh K Saini, Shahbaz Aman, Vijaykumar Muley, Gandhi Khaba Wairokpam, Zahoor Ahmad Parray, Anjuman Sahani, Amit Pathania

## Abstract

1.

Abiotic stresses like nitrogen deficiency and soil salinity are major factors contributing to low crop yields. The use of selective biofertilizers alleviates both types of stress. In this study, we investigated the biofertilizer activity and plant growth-promoting properties (PGP) of *Rhodococcus jialingiae* RS1 through cytosolic proteome remodelling. We cultured RS1 under two conditions, i) without and ii) with 6% NaCl, in nitrogen-deficient defined Burk’s medium. Under dual stress of nitrogen limitation and salt stress, Orbitrap LC-MS/MS proteomics revealed one-quarter of the proteome remodelling, particularly the upregulation of ribosomal synthesis and protein repair systems. As expected, we found high expression of EctC, an ectoine synthase, a key enzyme in osmolyte biosynthesis. Additionally, ribosomal and translational-associated factors, including RpsL, RpsS, RpsT, RpsR1, RplV, RplL, RplA, and elongation factor Tuf, were highly expressed, suggesting enhanced translational fidelity under dual stress. High levels of DNA protection protein, Dps suggest dual stress may lead to DNA damage. Upregulation of chaperones, environmental sensors (KinE), and redox transcriptional factors like WhiB3, Hsp18, AhpC, and MetE suggests protein misfolding and oxidative stress. Metabolic modulations were evident through high expression of IlvA, NAD-dependent glutamate dehydrogenase, lipid/envelope-remodelling enzymes, cutinase/esterases, lipases, endopeptidases like NlpC/P60 and transport systems. In contrast, proteins involved in urease structural components (urea-G), nitrogen regulators and ammonium transporters (GlnK and Amt) were downregulated. Dual stress may lead to an energy crisis, prompting strategic shifts away from high-ATP-dependent ureolytic nitrogen-scavenging pathways towards lower-energy nitrogen-assimilating routes, such as IlvA-mediated deamination and NAD-dependent glutamate dehydrogenation. Genetic manipulations of the above-mentioned genes or their homologues across the genera of microbes, plants, and crops may enhance resilience to abiotic stresses. Our studies reveal stress-responsive genes and biochemical pathways that could be used to improve transgenic efficacy in nitrogen-limited, saline soil and other (a)biotic stresses.

**Global Proteome Profiling *of Rhodococcus jialingiae* RS1 to Develop Transgenics:** 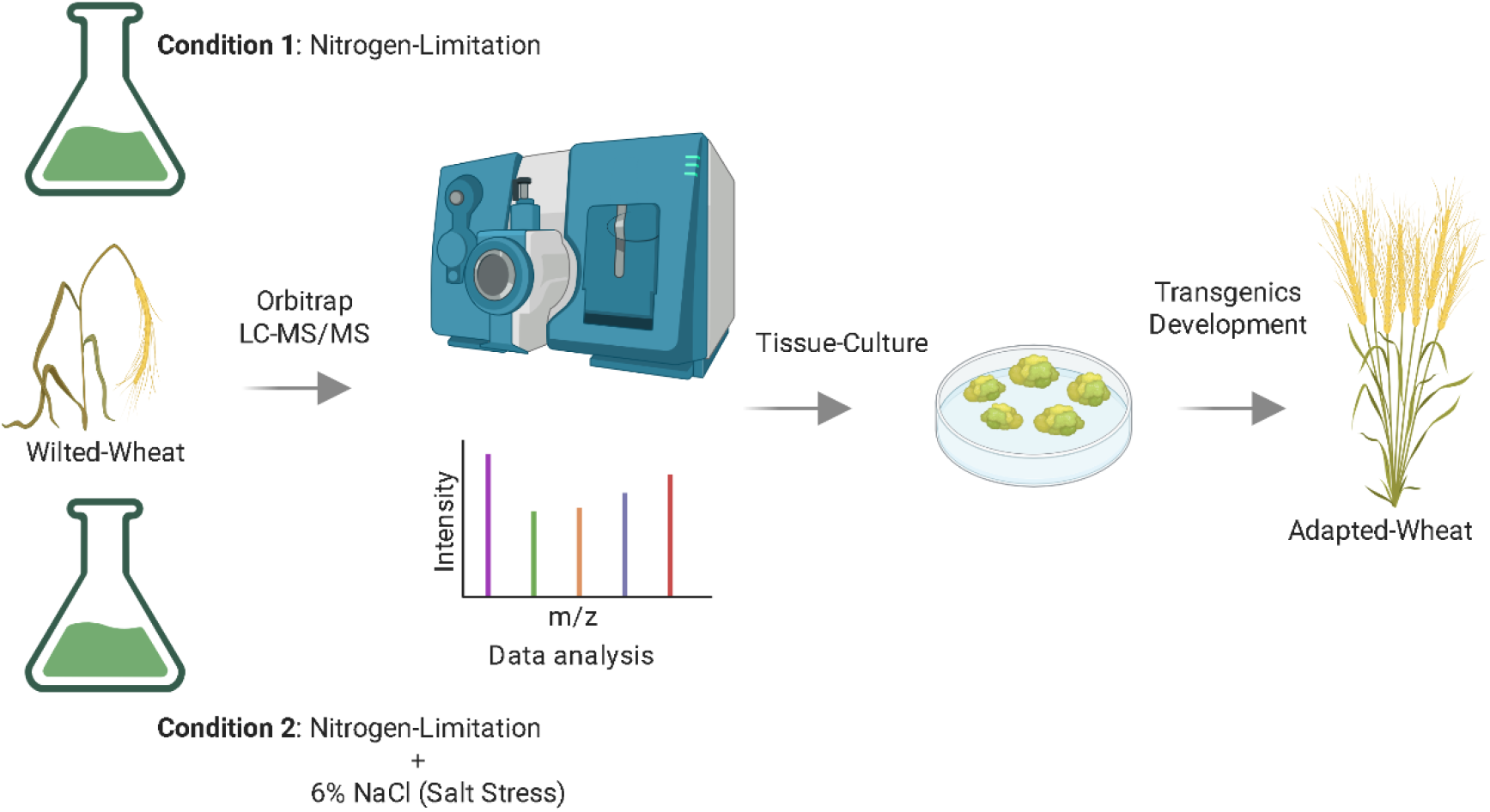

## 2. INTRODUCTION

Worldwide, overexploitation and poor farming practices are increasing salinity and decreasing nutrient levels, particularly nitrogen in agricultural soils, hindering crop productivity (1). In agronomy, bacteria that enhance nutrient availability through mechanisms, like nitrogen fixation, phosphorus solubilisation and potassium uptake, are commonly classified as biofertilizers (2–4), whereas bacteria that produce osmolytes (5,6), phytohormones and maintain stress hormone balance (7), and overall facilitating plant/crop growth (8), are referred to as PGPB (Plant growth-promoting bacteria). Within this framework, the genus *Rhodococcus*, a member of the actinobacteria, is widely utilised as a bioremediation agent (9) and thrives in various soil types, including those with high and low pH levels and high salt concentrations(7).Under high salinity, *Rhodococcus* accumulates various osmolytes and lipids to maintain osmotic balance and protect proteins from denaturation (10), while in parallel producing phytohormones and enhancing overall plant growth, thereby fulfilling the criteria of a PGPB (11)(12). *Rhodococcus* species possess the genetic machinery necessary for nitrogen fixation (*nif* and *rnf* genes), enabling them to act as biofertilizers by providing plants with essential nitrogen nutrients (13,14). This suggests that cohabitation of *Rhodococcus* with crops/plants in the same soil can promote their growth and improve the soil quality. To better understand the adaptive characteristics of microbes in simulated plant-associated environments, a high-throughput, comprehensive systems-level approach, protein profiling (proteomics) is utilised. Protein profiling captures changes in the global proteome in response to environmental stimuli and gives a glimpse of what changes happen in real time at molecular and biochemical pathways (15). This is exemplified by diazotrophs, nitrogen-fixing bacteria, where the bacterial proteome shifts towards nitrogen assimilation during nitrogen deficiency, whereas halotolerant PGPB engage in osmo-protectant mechanisms, like production of compatible solutes and osmolytes (18,19). This distinct profile distinguishes the nitrogen-producing bacteria (biofertilizer) like *Crocosphaera watsonii, Anabaena 7120*, and *Geomonas sp*(18–20). from halotolerant (PGPB) like *Bacillus subtilis, Pseudomonas pseudoalcaligenes*, and *Exiguobacterium oxidotolerans* (21). In addition, bacteria like *Pseudomonas stutzeri* and *Klebsiella pneumonia*, which serve both the purposes of nitrogen fixation and saline tolerance and live near the roots of plants/crops (rhizobacteria) have been shown to improve the rice crop yield (22).

In this study, we examined the proteome remodelling of the RS1 strain under nitrogen-limiting and salt-stress conditions in vitro, with the aim of identifying key proteome signatures important for RS1 adaptation and their potential use in future transgenics. Under dual stress, RS1 upregulates ribosomal proteins, osmo-protectant genes, and biosynthesis of oxidoreductases, while repressing many proteins, especially involved in urease synthesis. Protein–protein interaction analysis shows that RS1 acclimates by maintaining high translational fidelity, rerouting energy to less demanding pathways, and upregulating proteins that rescue core components of the central dogma-DNA, RNA and proteins.

## 3. MATERIALS AND METHODS

### Bacterial strain and growth conditions

*Rhodocccus jialingiae* strain 31455 was procured from ATCC (23). The strain was named RS1 (*Rhodococcus jialingiae* RS1). RS1 glycerol stocks were stored in -80 °C. Glycerol stocks were streaked on Nutrient agar plates (HiMedia, India) and incubated at 30ºC for 48 hours. A single colony was inoculated in 10 ml of Nutrient broth and incubated overnight at 30 ºC at 180 RPM. As reported earlier (13), the overnight-grown culture reached the mid-log phase. The overnight culture was subcultured at a 1:1000 dilution into 300 mL of nutrient broth. The flasks were incubated at 30 ºC /180 RPM until the OD600 reached 0.6. The cells were pelleted, washed twice with sterile 1X PBS, and resuspended in either 300 mL of nitrogen-deficient Burk’s medium (HiMedia, India) as a control (RS1N) and the same volume of Burk’s media +6% NaCl as a test (RS1NS). The flasks were further incubated at 30 ºC and 180 RPM for 6 hours. Post incubation, the bacterial cells were harvested by pelleting at 4000 RPM for 10 minutes at RT. These pellets were stored at -80 ºC until further use. The experiment was done with 2 independent biological repeats.

### Cytosolic protein extraction

RS1 cell pellet (∼1g) of two conditions, i) RS1N and ii) RS1NS, which were stored in -80ºC, were thawed and dissolved in 750 μL of lysis buffer (containing 7 M urea, 2 M thiourea, 4% CHAPS, 1× PMSF protease inhibitor), and 0.5 gram of 1 mm-sized zirconium glass beads were added. The samples were incubated at room temperature for 3-4 hours to weaken the hard cell walls. The samples were transferred to the Neuation bead beater/tissue homogeniser (iRupt 24, India). Parameters were set to 10 cycles at 4500 RPM, with each cycle lasting 1 minute. The cell lysates were centrifuged at 13000 RPM for 30 minutes. The Supernatant was collected to obtain cytosolic proteins.

### Protein precipitation and quantification

150 µL of each of the above-mentioned prepared samples was mixed with 1.4 mL of acetone and incubated overnight at -80 °C. The samples were centrifuged at 13K RPM for 30 minutes at 4 °C, and the supernatant was discarded. 1 mL of 80% ethanol was added, and the mixture was centrifuged at 4 °C for an additional 30 minutes at 13K RPM at 4 °C. The supernatant from all samples was discarded, and the pellets were dried at 37 °C to remove any excess ethanol and water. Finally, the dried pellets were dissolved in 50 µl of a solution containing 4 M guanidinium HCl and 50 mM Tris. The Pierce BCA protein assay kit (Thermo Scientific, USA) was used to quantify protein.

### Orbitrap-derived LC-MS/MS (Liquid Chromatography-tandem Mass Spectrometry) analysis

#### In-Solution Trypsin Digestion

Protein samples at a concentration of 1 µg/µl were processed further at the Central Instrumentation Facility (CIF), University of Delhi South Campus, for LC-MS/MS analysis using an Orbitrap (ThermoFisher Scientific, USA). Protein samples undergo the standard protocol used for the in-solution digestion method. In short, the protein samples were further treated to yield a solution containing 6 M guanidium HCl along with 0.1 M Tris (pH 8.0). This denaturing buffer concentration is necessary because it disrupts the protein’s 3D structure and solubilises it. DTT (dithiothreitol) of concentration 20 mM was added to reduce protein samples, and sample mixtures were incubated for 1 hour at room temperature. Next, DTT was removed, and 40 mM IAA (iodoacetamide) was added to the sample mixture for alkylation. Following this, the samples are incubated for 30 minutes in the dark. The DTT was used to stop the reaction. The trypsin was then added at a 1:20 to 1:100 ratio (w/w), and the samples were incubated overnight at 37 °C for complete proteolysis. The digested sample peptides were stored at -20 °C until their use.

#### Peptide Desalting

With the help of Thermo Scientific™ Pierce™ Peptide Desalting Spin Columns (Cat. No. 89851/89852), desalting of peptides was performed as per the protocol of the manufacturer. Peptides were reconstituted in 0.1% formic acid and processed further with LC-MS/MS analysis through Orbitrap (ThermoFisher Scientific, USA).

#### LC-MS/MS analysis

It was performed using a Thermo Scientific Easy-nLC 1200 system coupled to a Q Exactive Orbitrap mass spectrometer. Approximately 2 µg of peptide was loaded onto a PepMap RSLC C18 column (2 µm particle size, 75 µm × 50 cm, Thermo Scientific), maintained at 60°C. The mobile phases consisted of solvent A (98% water, 2% acetonitrile, 0.1% formic acid) and solvent B (20% water, 80% acetonitrile, 0.1% formic acid). A linear gradient was applied over a total run time of 120 minutes with the following settings: 0% B for 5 minutes, gradually increasing to 30% B over 100 minutes, followed by a rapid rise to 95% B for column washing and then re-equilibration to 0% B. The flow rate was maintained at 300 nL/min. All solvents used were of LC-MS grade (Fisher Scientific), and a lock mass of 445.12003 Da was used for internal calibration. The Q Exactive Orbitrap operated in positive ionisation mode with a total run time of 120 minutes. MS1 scans were acquired at a resolution of 70,000 with a scan range of 350–2000 m/z, automatic gain control (AGC) set to 3e6, and a maximum injection time (IT) of 50 ms. Data-dependent MS2 acquisition was performed with the top 10 most intense precursor ions selected for fragmentation using higher-energy collisional dissociation (HCD) at a normalised collision energy of 27. MS2 scans were acquired at a resolution of 17,500 with AGC set to 1e5 and IT of 120 ms. The isolation window was set to 1.5 m/z with a dynamic exclusion time of 50 seconds. The fixed first mass was set to 100 m/z.

### Proteomic data processing

Raw files were processed in Proteome Discoverer 2.4 (Thermo Scientific) against UniProt databases for *Rhodococcus erythropolis*. Search parameters: 10 ppm precursor tolerance, 0.02 Da fragment tolerance, trypsin (2 missed cleavages), fixed carbamidomethylation (Cys), 1% FDR (PSM/protein level). Label-free quantification used Minora feature detection.

### Scatter plot analysis of RS1N and RS1NS replicate

To validate the consistency of the biological duplicates for RS1N (Nitrogen-deficient Burk’s medium) and RS1NS (Burk’s medium + 6% NaCl), a scatter plot was used. The log2 abundance ratios of proteins across replicates (RS1N & RS1NS) were provided, and the duplicates quantified in both were selected and uploaded to R (version 4.5.0). R GUI generates a pairwise scatter plot with one repeat on the X axis and another repeat on the Y axis. Here, the Pearson correlation coefficient (r) was determined by Pearson Cor(24). The generated scatter plot was exported as high-resolution images and displayed to prove the consistency of the biological repeat.

### Venn Diagram

Web-based Venny 2.1 tool was used to analyse and visualise the distribution of proteins in two conditions, i) RS1N and ii) RS1NS. The protein accession numbers from each biological repeat (RS1N1 Vs RS1NS1 and RS1N2 Vs RS1NS2) were used (Supplementary Files X3, X4, X5 and X6). FDR (false discovery rate) was filtered at a q-value ≤ 0.05. The accession number of the proteins belonging to the biological repeat pair (RS1N1 vs RS1NS1; RS1N2 vs RS1NS2) was pasted as a two-input comparison into the web-based Venny 2.1 tool (25). The software analyses the number and percentage of shared proteins in each and between the two conditions, and the results are displayed in a high-resolution image.

### Heatmap

For the label-free LC–MS/MS cytosolic proteomic dataset of two conditions, i) RS1N and ii) RS1NS, a total of 1215 proteins with reliable quantitative values were obtained after initial filtering and basic missing-value. For heatmap visualisation, proteins were first restricted to those showing statistically significant differential abundance between RS1NS and RS1N (adjusted p-value ≤ 0.05). Among these significant proteins, the 50 most strongly regulated were selected based on the largest absolute log2 fold change, ensuring inclusion of both highly upregulated and highly downregulated proteins. For the selected proteins, raw intensities in the four samples (RS1N1, RS1N2, RS1NS1, RS1NS2) were transformed as log10 (abundance +1) and then scaled on a row-wise basis to z-scores, so that each protein’s relative change between conditions was emphasised independent of its absolute intensity. The resulting matrix was visualised as a hierarchical clustering heatmap using the pheatmap package (R 4.5.0), with Euclidean distance and complete linkage applied to proteins (rows) and the columns ordered by biological condition (RS1NS vs RS1N). In the final heatmap, higher-than-average abundance for a given protein (relative to its own mean) is displayed in red, and lower-than-average abundance is displayed in blue, highlighting the core set of significantly reprogrammed proteins under salt stress.

### Volcano Plot

Protein expression differences of two conditions, i) RS1N and ii) RS1NS, were analysed in R (version 4.5.0). For each protein, log2 fold change (log2FC) and Benjamini–Hochberg false discovery rate (FDR)-adjusted p-values (padj) were calculated to control for multiple testing. Proteins with padj ≤ 0.05 and log2FC ≥ 1 were classified as upregulated, whereas proteins with padj ≤ 0.05 and log2FC ≤ −1 were classified as downregulated; all remaining proteins were considered non-significant. Volcano plots of log2FC versus −log10 (padj) were generated using a white background and coloured points (red, upregulated; blue, downregulated; grey, non-significant) to visualise global expression changes.

### Gene Ontology Enrichment and Network Analysis

Differentially expressed cytosolic proteins in two conditions, i) RS1N and ii) RS1NS, were divided into upregulated and downregulated sets. The two sets were analysed for Gene Ontology enrichment using ShinyGO v0.85 across three domains: Biological Process, Cellular Component, and Molecular Function (Supplementary File X10 and X11). The FDR cutoff, 0.05 was set for all three domains, and the annotated proteome of *Rhodococcus erythropolis*, was used as the background. Significant terms were identified using hypergeometric over-representation with Benjamini–Hochberg FDR correction and visualised as dot plots ranked by fold enrichment. Proteins from enriched terms in each GO domain were combined and submitted to STRING (*Rhodococcus erythropolis* reference, confidence ≥ 0.40, no additional interactors); the resulting interaction networks were clustered using the Markov Cluster Algorithm (MCL, default inflation), and highly connected nodes within clusters were considered hub proteins for functional interpretation.

## 3. RESULTS

### A Pearson Correlation Coefficient (r), exceeding 0.75 in 2 independent protein profiles

We first tested the growth of *Rhodococcus jialingiae* RS1 in the presence of Burk’s medium and Burk’s medium with varying NaCl concentrations (4% to 10%) (26). We observed that a 6% NaCl concentration effectively induced short-term salt stress, as RS1 growth remained nearly static under this condition (Supp FIG 1 and Supplementary File X0). Based on this, we performed a label-free quantitative cytosolic global proteomic analysis for RS1 under two different conditions: i) Nitrogen-deficient (RS1N) and ii) Nitrogen-deficient supplemented with 6% NaCl (RS1NS). Each condition was analysed using two biological replicates.

**FIG 1.**
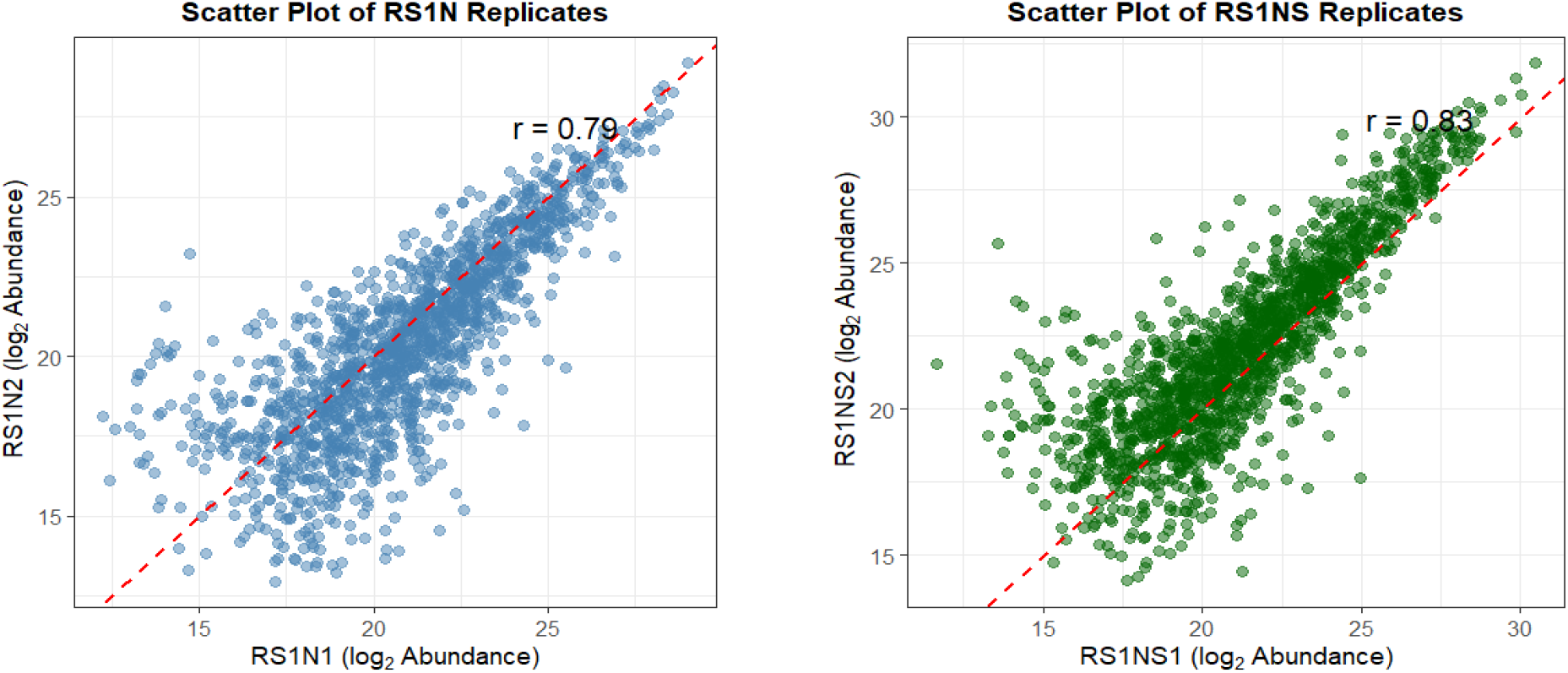
Scatter plot analysis of the Log2 abundance values of proteins. R (version 4.5.0) was used to generate a scatter plot of the biological replicates using Log2 of protein abundance values (27). **(A)**, The X-axis represents protein samples from RS1N1, and the Y-axis represents protein samples from RS1N2. In RS1N1 vs RS1N2, the straight dotted red line represents the correlation between two replicates (r=0.79). A blue dot represents the protein abundance of a biological replicate. **(B)**, Here, notions are the same as in (A), and the replicates are shown as RS1NS1 vs RS1NS2 (r=0.83).

biological replicates. To verify reproducibility, a scatter plot analysis was performed across independent biological repeats (27). For RS1N biological repeats (RS1N1 and RS1N2), the Pearson Correlation Coefficient (r) is 0.79, while in RS1NS repeats (RS1NS1 and RS1NS2), the r value is 0.83 (Fig. 1, A and B). These r-values confirm that, on average, biological repeats represent the same set of expressed protein(28). The details of RS1N1, RS1N2, RS1NS1, and RS1NS2 are given in the supplementary files X1 and X2.

### Dual stress causes a quarter (∼25%) change in the proteome

To make statistically valid conclusions, filtered proteins with their abundance values (q value ≤ 0.05; for details, see Materials and Methods and Supplementary Files X3, X4, X5 and X6) were compared across two conditions (RS1N and RS1NS) in 2 replicates (RS1N1, RS1N2, and RS1NS1, RS1NS2). As shown in Figure 2, around 25% proteins were specific to each condition (RS1N1-22.8 %; RS1N2-27.6% and RS1NS1-28.3%; RS1NS2-25.7%), and approximately 50% proteins were expressed in both conditions (Figure 2, A, overlapped region, 48.9% and B, overlapped region, 46.7%). The data support the consistency of our scatter plot analysis (Fig. 1). Approximately 25% of condition-specific proteins in RS1N or RS1NS would be selected for further study.

**FIG 2.**
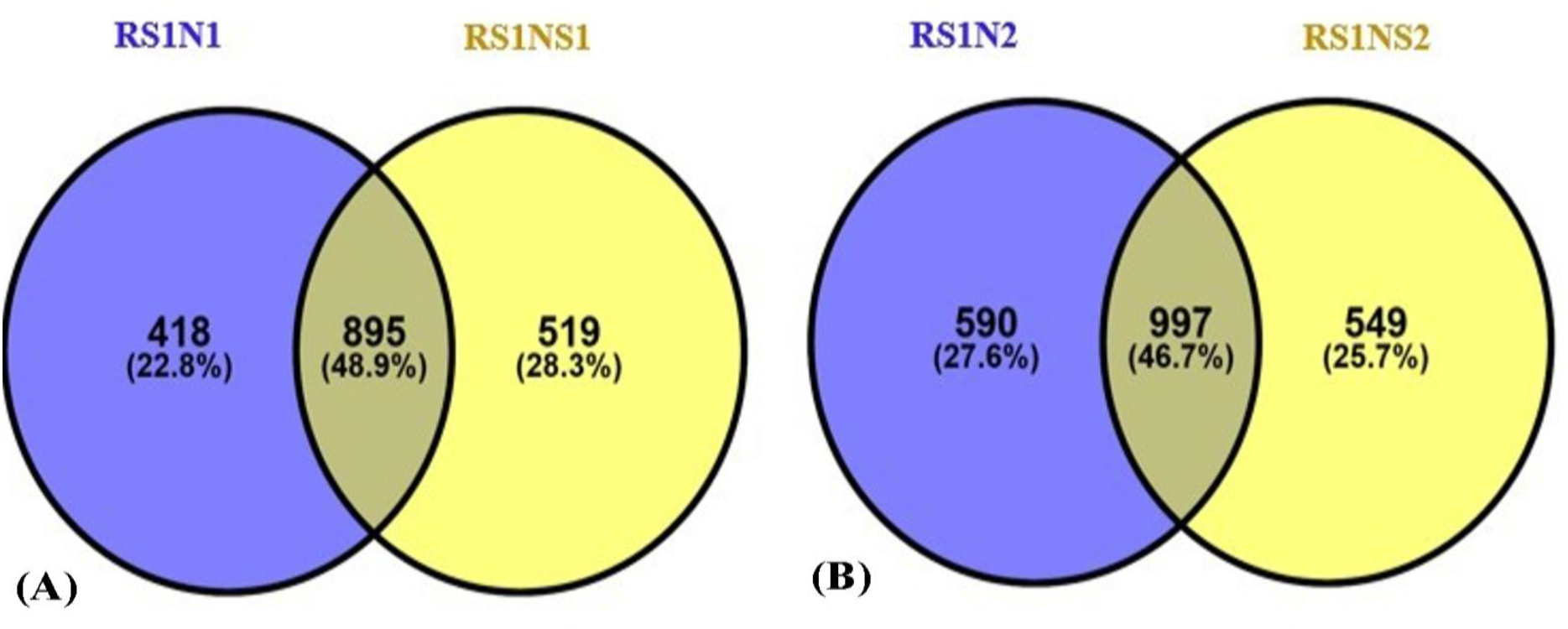
Overlapping protein profiles in replicates of RS1N (Nitrogen-deficient) and RS1NS (Nitrogen deficient+6% NaCl). The Venny 2.1 tool was used to generate Venn diagrams comparing protein profiles in (A) RS1N1 vs RS1NS1 and **(B)** RS1N2 vs RS1NS2. The actual number of proteins (from left to right, 418, 895, 519, equal to 1832 (100%) in the 1^st^ replicate, and 590, 997, 549, equal to 2136 (100%) in the 2^nd^ replicate), and their percentages, are given in each case. Overlapping regions of Vienn Diagrams represent proteins which are expressed in both conditions, without and with 6% NaCl. The protein profiles of RS1N replicates are represented in blue, while those of RS1NS replicates are shown in yellow.

### Dual stress’ Heatmap Signatures: Ribosomal proteins and Oxido-reductases

Heatmap of proteins, 50 in number, which are either highly expressed/repressed in dual stress (RS1NS over RS1N) is given in Figure 3 (For details, see Materials and Methods, and Supplementary file X7). In RS1NS condition, one group of proteins, which are in abundance belong to ribosomal proteins such as rpsZ, rpsR, rpmF and rpmI which are seems reported to participate in stress adaptation (29,30). There are many proteins related to stress adaptation, macromolecule biosynthesis, including cell-envelope and lipid-metabolism enzymes such as cutinase/esterase-C0ZYC0, antibiotic-biosynthesis monooxygenase-A0A0C3A5T6, acetyl-CoA C-acetyltransferase-A0AAX3V6P9, 2-oxoisovalerate dehydrogenase-A0A6G9D385 and DNA-protection protein-Dps-A0A0C3AD17 are upregulated. Some uncharacterized but strongly up-regulated proteins, such as A0A5P3G4V6, E9RG82, C1A1R8, A0A8I1A801, A0AAX3V3J8, A0A0C3A4E7, A0A8I1D6I1, A0A0C3A3W1, likely participate in halotolerance (31). Here, Acetyl-CoA C-acetyltransferase-A0AAX3V6P9 is involved in the mevalonate pathway for secondary metabolite production and the synthesis of triacylglycerols, which may lead to osmotic balance and cell envelope modulations (32). This indicates that under RS1NS, the bacterium’s first priority is translational fidelity as reported previously (37,38). It seems bacteria direct their metabolic flow into branched-chain and storage lipids, remodel their cell walls, and reinforce genome protection and other specialised stress-response functions to withstand dual pressure (33). Conversely, several proteins that are down-regulated in RS1NS over RS1n are closely associated with nitrogen scavenging, oxidative-stress defence and membrane/lipid turnover. Such as UreC, the nitrogen-regulatory PII protein, GlnK and the ammonium transporter-A0A5N5EA30, as well as the catalase-peroxidase KatG and multiple enzymes linked to lipid and cell-wall dynamics (lipase-A0A6G9CM33, NlpC/P60 endopeptidase-A0AAX3V130, MBL-fold hydrolase-A0A8I1D6E2, NTF2-family protein-A0A0C2WDA0 and the transport proteins UgpC and A0A8I1A3H6). This suggests that, by downshifting nitrogen scavenging and oxidoreductases, bacteria are allocating their energy to translational-fidelity and osmoregulation.

**FIG 3.**
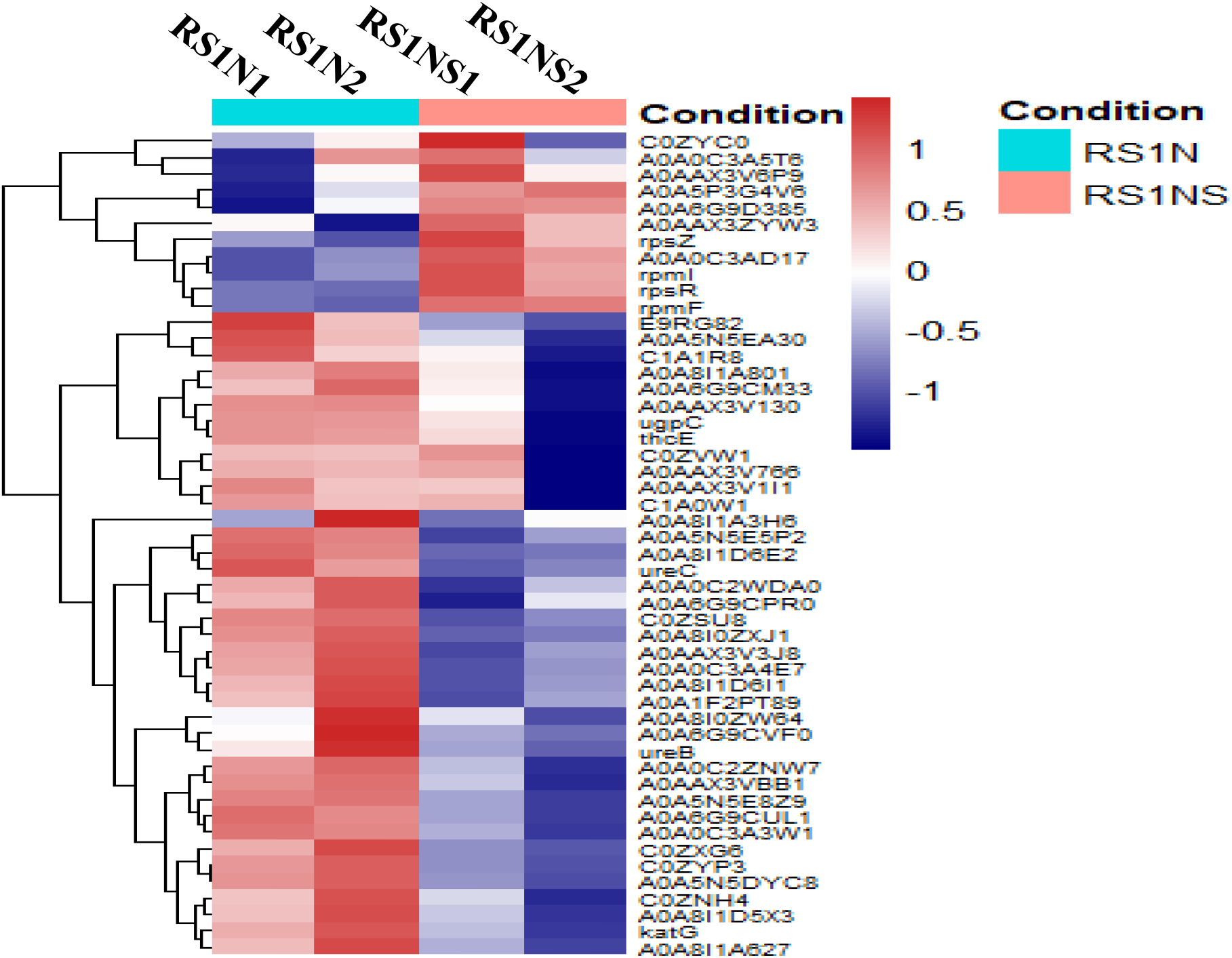
Heatmap of specific proteins in growth conditions without and with 6% NaCl. R (version 4.5.0) was used to generate a Heatmap (34) of 2 biological replicates: Nitrogen deficient (RS1N-Turquoise Blue); RS1N1, RS1N2, and Nitrogen deficient with 6% NaCl (RS1NS-Carrot Red); RS1NS1, RS1NS2). The first 50 highly regulated proteins are shown here. Shades of red (0 to 1) and shades of blue (0 to -1) represent proteins which get highly expressed and repressed. Many genes are not annotated, hence are presented with their accession numbers.

### Dual stress adaptation involves 2 components: Minimal maintenance and reducing the energy-consuming resources

Life in all organisms is sustained through an active Central Dogma (35). The central dogma is the flow of information from DNA to RNA and from RNA to protein (36). Stress hampers the normal operation of the central dogma, and the organism’s cellular machinery responds to stress, addressing it and restoring the central dogma’s activity. Meanwhile, to remain biologically active and sustain its central dogma, minimally important proteins are highly expressed (37,38). Along with this, the energy requirement would be maintained by shutting down activities that have not been fruitful for sustenance or for restoring the normal central dogma(39). In our Volcano’s plot (Figure 4 and supplementary file X8), ribosomal proteins like RplF, RplR, RplU, RplV, RpmA, RpmB, RpmF, RpmI, RpmJ, RpsS, RpsL, RpsR, RpsT and RpsZ, needed for sustaining translation activities, are highly expressed (29). In dual stress, protein foldings will be affected, and hence, chaperones like heat shock protein, Hsp18, which may help in protein foldings, are highly expressed (40,41). Many types of abiotic stress, in our case high salt and nitrogen deficiency, lead to the formation of ROS (Reactive Oxygen Species), which ultimately cause oxidative stress (42,43). Hence, to adapt to it, enzymes like alkyl hydroperoxide reductase, AhpC, get highly expressed (44,45). Methionine is a more susceptible amino acid to oxidative stress in the presence of ROS (46,47). Hence, to get its supply, the expression of methionine synthesis, MetE, increased here. Osmo-protection is supported by the up-regulation of ectoine synthase, EctC (48,49). For NH4^+^ production, threonine dehydratase (encoded by *ilvA*) expression is increased, it deaminates threonine and NH4^+^ get produced (50). WhiB expression normally gets turned on during stress management like antibiotic resistance and in our case for salt stress in nitrogen deficient medium (51–53). Dxr, dihydroorotate dehydrogenase, high expression leads to isoprenoid synthesis, which helps in antibiotic and metabolic stress (54–56).

**FIG 4.**
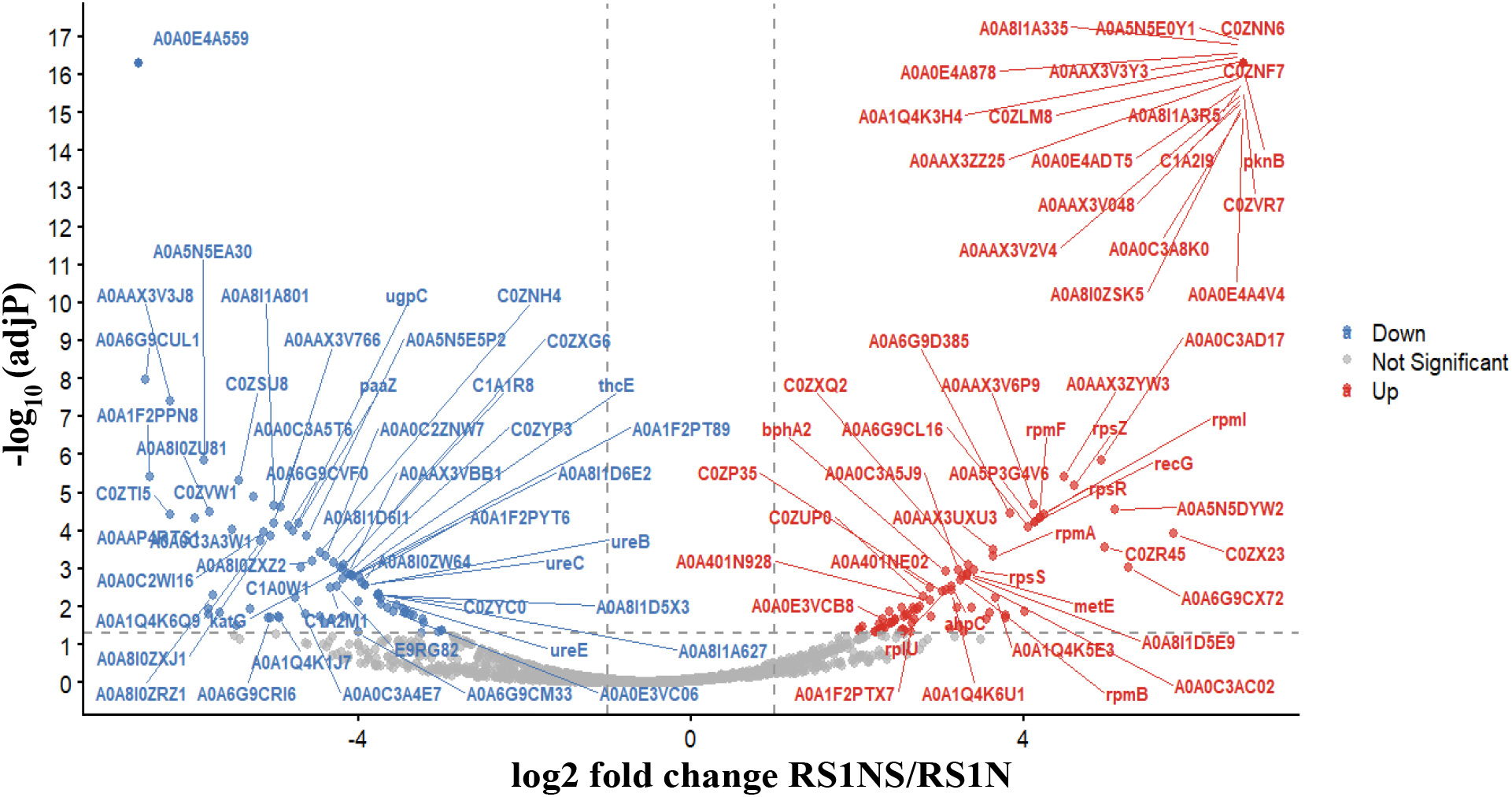
Volcano plot of differentially abundant cytosolic proteins in dual stress. The X-axis shows the log2 fold change in protein abundance (RS1NS; dual stress-nitrogen deficient+6%NaCl over RS1N; nitrogen deficient), and the Y-axis shows the −log10 adjusted p-value for each protein. Each point represents one or more protein-coding genes. Proteins with log2 fold change ≥ 1 and adjusted p-value ≤ 0.05 are highlighted in red and correspond to significantly up-regulated proteins in RS1NS, whereas proteins with log2 fold change ≤ −1 and adjusted p-value ≤ 0.05 are highlighted in blue and correspond to significantly down-regulated proteins in RS1NS (for details, see Materials and Methods). Grey points denote proteins that do not meet these thresholds and are considered not significantly changed under the chosen criteria.

To reorganize the energy or resources, *Rhodococcus jialingiae* RS1 represses structural and accessory urease components (UreA, UreB, UreC, UreE and UreG). It emphasises that if NH4^+^ is supplemented with IlvA high expression, other sources of it is blocked or get repressed. Central-metabolic enzymes used in glucogen metabolism GlgP and cell wall synthesis, UDP-N-acetylglucosamine pyrophosphorylase, GlmU are also strongly decreased, suggesting reduced glycogen mobilization and peptidoglycan precursor synthesis relative to control conditions. Additional down-regulated proteins include the mechanosensitive channel MscL, transport protein UgpC, catalase KatG and several other oxidoreductases and hypothetical proteins (e.g. A0A6G9CUL1, A0A0C2WDA0, A0A8I1A4Z7, A0A6G9CWB3), highlighting an energy shift away from these pathways (for details see Supplementary file X7).

### Ribosome–cell envelope crosstalk and repression of urea catabolism lead the biological pathways

ShinyGO biological process analysis as shown in Fig. 5, exhibits strong differentiation in biological pathways. The proteins of the ‘Cellular Macromolecule biosynthetic process’ pathway, get around 6.0-fold enrichment in RS1NS (Nitrogen deficient+6% NaCl) over RS1N (Nitrogen deficient) (for details, see the materials and methods, Supplementary File X9 and X10). This enrichment is mainly driven by the ribosomal proteins, as shown in FIG 5A and Supplementary File X9, and associated biosynthetic factors such as AKD95549.1 and AKD97324.1. In addition, other pathways like, translation, peptide metabolic/biosynthetic processes, macromolecule and amide biosynthesis, cellular macromolecule metabolism were also enriched (supplementary file 9). Ribosomes/protein factories are the basic unit of translation (57) . To keep protein factories alive in nitrogen-deficient and 6% NaCl, proteins involved in the assembly and stabilisation of 30S and 50S ribosomes (70S) are highly expressed. Here, RpsL, RpsS, RpsT, and RpsR1 encode 30S ribosomal proteins, and RplV (L22) supports 50S assembly. On the other hand, enriched pathways, in the repressed protein profile during the RS1NS condition, are related to nitrogen metabolism (Fig. 5C).

**FIG 5.**
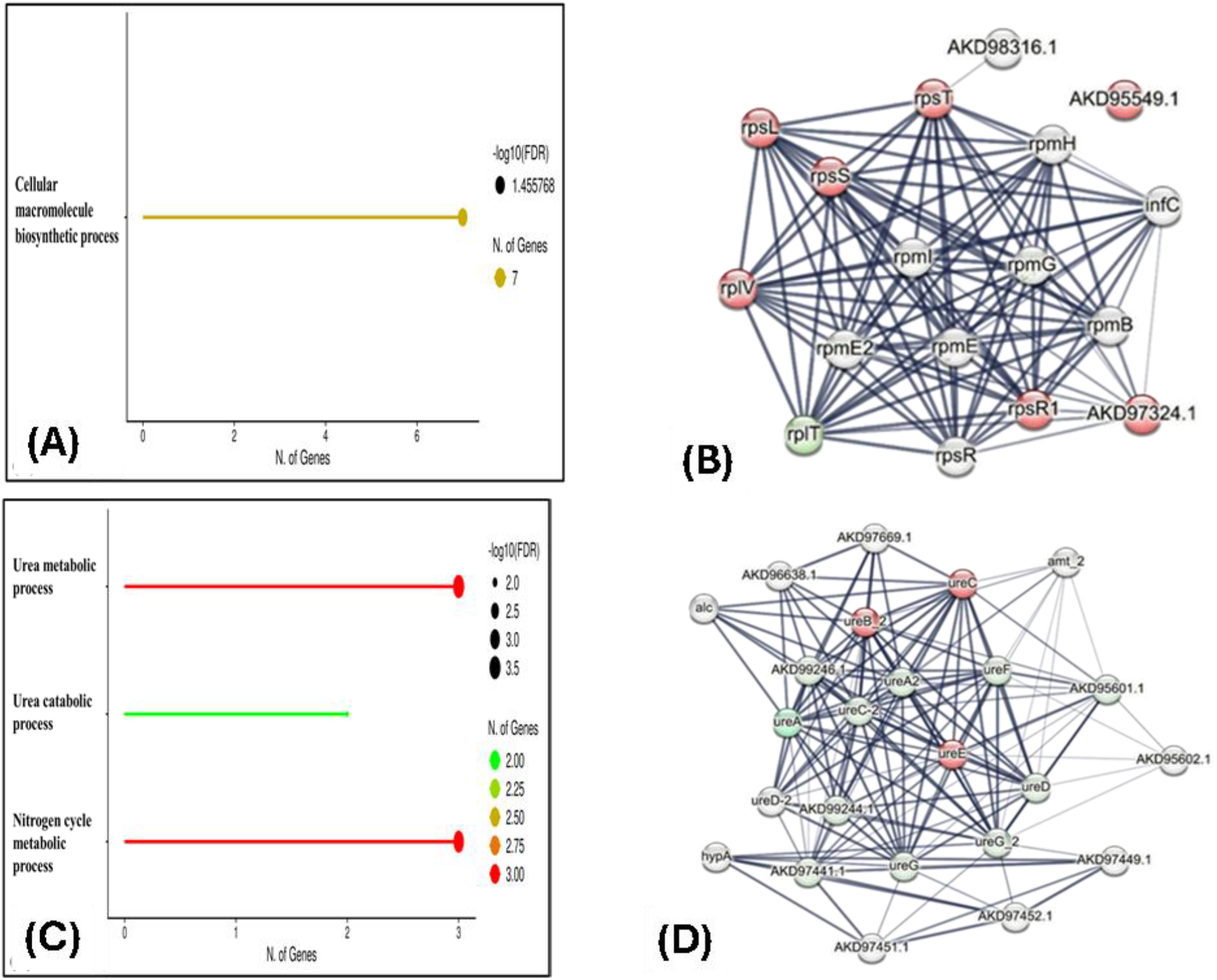
Enrichment analysis of biological pathways and their PPI (Protein-protein interaction) networks. ShinyGO 0.85.1 is used to analyse enriched pathways (58) as shown in A and C. STRING version 12.0 is used for protein-protein interaction (59) as shown in B and D. **A and C** Here, the input comes from the proteins identified in LC-MS/MS analysis. The X axis indicates the number of gene each ShinyGO pathway possess, and the Y axis shows the name of the ShinyGO pathway. The graph in **(A)** shows an enriched pathway in which genes encoding ribosomal proteins are highly expressed, involved in protein/macromolecule synthesis. Proteins of the macromolecular synthesis pathway, represented as ‘Cellular Macromolecule biosynthetic process’, have been shown on the Y-axis. In (C), repressed proteins of dual stress have been used as input and 3 enriched pathways, all related to the Nitrogen metabolism are shown. **B and D)** proteins that give rise to graph **A**, have been used as input for **B** and proteins that appear in graph **C** have been used as input for **D**. Nodes in red represent the proteins used as input for the STRING analysis. The grey nodes represent proteins that interact with the input proteins. Edges represent the interaction and intensity of the edge colour, grey to dark represents the confidence of the interaction.

For protein-protein interaction (PPI) through STRING (60) as shown in Fig. 5B, the input data is taken from Fig. 5A. On the same line, AKD97324.1 is supposed to be involved in ribosomal RNA modulation. STRING highlights the interaction between ribosomal proteins and cell envelope assembly. This notion is supported by the high expression of the protein AKD95549.1, which is involved in phospholipid and lipid carrier turnover (part of the cell envelope). In addition to ribosomal proteins, another protein in the red node is AKD97324.1, supposed to be involved in RNA modulation. The white and pale-colored nodes/proteins which appear only in outputs of the STRING and not due to ShinyGO are infC, rplT, rpmB, rpmE, rpmE2, rpmG, rpmH, rpsR, AKD98316.1, etc. These proteins appear to be finely associated with other ribosomal proteins. *infC*, an essential gene that encodes IF-3 (Initiation Factor), helps form the translational initiation complex. MCL (Markov Clustering) (Supplementary file X12) is also a part of STRING analysis, reinforcing the above ideas. It indicates that under nitrogen-deficient and 6% NaCl conditions, there is an interconnection among ribosome biogenesis, cell envelope synthesis, and the efficiency of the translational initiation complex. It appears that maintaining minimal function of the central dogma (DNA to RNA and RNA to Protein) is the primary criterion for adaptation in salt- and nitrogen-deficient conditions

On the other hand, enriched pathways in the repressed protein profile under the RS1NS condition are related to nitrogen metabolism (Fig. 5C, (for details, Supplementary File X11 and X14). Three repressed enriched pathways are i) urea metabolic process, ii) nitrogen cycle metabolic process, and iii) urea catabolic process, supported by ureB_2, ureC and ureE. As shown in Figure 5D, the input data is taken from Figure 5C, for the STRING network (for details see Materials and Methods, and Supplementary X14). STRING network analysis of individual ureolytic proteins revealed that these proteins are part of a larger, evolutionarily conserved urea nitrogen scavenging module made up of duplicate Urease subunits (ureA/ureA2, ureB_2, ureC/ureC 2), maturation factors (ureD/ureD 2, ureF, ureG/ureG_2, HypA/HypB/HypC/HupD) and a urea transporter (AKD95601. 1), a urea amidolyase (AKD97669. 1), allantoicase (alc) and a putative ammonium transporter (amt_2). It suggests that high-capacity urea hydrolysing protein modules is co-downregulated under Nitrogen deficient and 6% NaCl. Markov Cluster Algorithm (MCL) (Supplementary File X13) partitioning of the STRING network consistently defined these proteins as a single, densely connected cluster containing urease (an enzyme that hydrolyses urea into NH3 and CO2) structural subunits, which scavenge nitrogen from excretory material such as urea(61,62).

### Highly expressed proteins localised in ribosomes and the cell envelope

ShinyGO Cellular component enrichment analysis as shown in Fig. 6A, reveals strong gene expression across specific cellular compartments (for details, see the materials and methods, Supplementary File X15). The compartments like ‘Small ribosomal unit’ and ‘Ribosomal unit’ do belong to ribosomes, while the proteins enriched in the ‘external encapsulating structure’ and ‘cell wall’ terms, the ParA-type partitioning ATPase-Soj_3 and the PldB-type lysophospholipase/αβ-hydrolase-AKD97402.1 are key envelope-associated factors in *Rhodococcus*. Soj_3 supports the faithful inheritance of stress-adaptation loci, whereas AKD97402.1 contributes to phospholipid turnover and remodelling within the capsule-like cell envelope. It suggests that dual stress leads to the formation of proteins which improve ribosome biogenesis, translational fidelity and the cell envelope (63,64).

**FIG 6.**
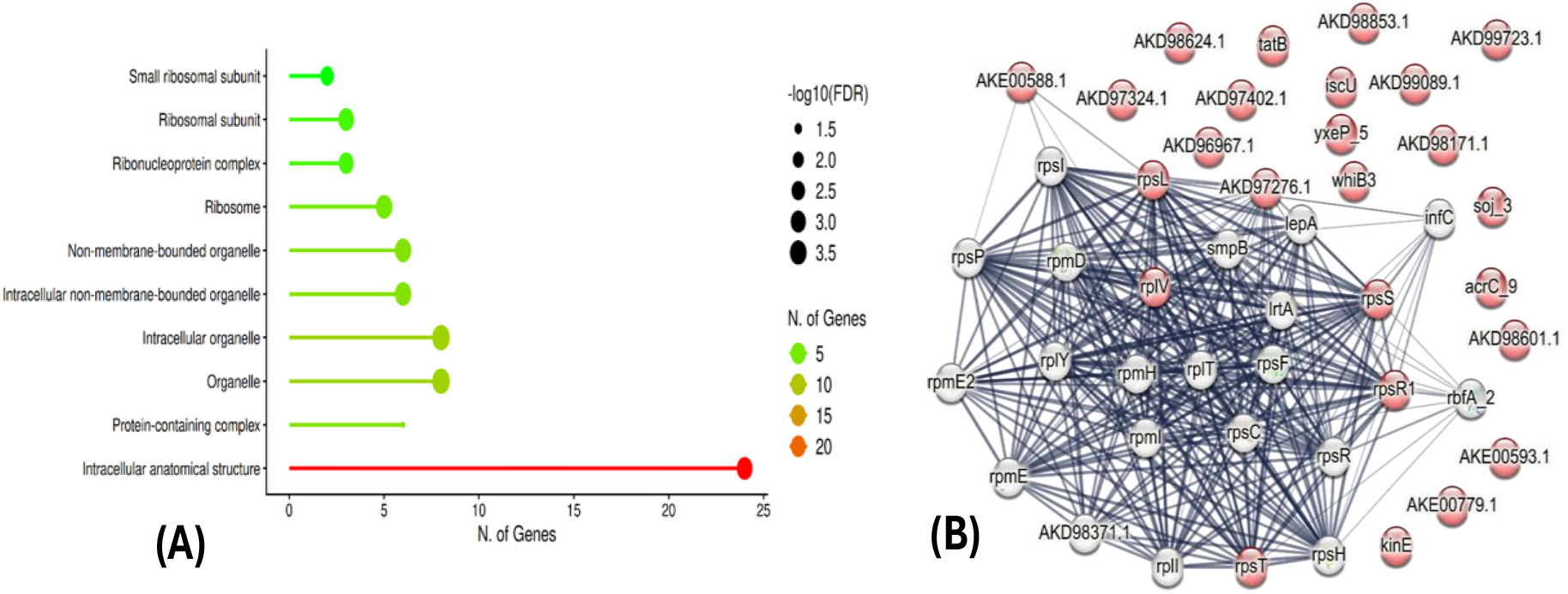
Cellular component enrichment of upregulated cytosolic proteins and their PPI (Protein-protein interaction) networks. ShinyGO 0.85.1 is used to analyse cellular components(58) as shown in A. STRING version 12.0 is used for protein-protein interaction(59) as shown in B. **A )** Here, the input comes from the proteins and accession numbers which are upregulated in RS1NS condition. The X axis indicates the number of gene each ShinyGO pathway possess, and the Y axis shows the name of the ShinyGO pathway. The graph in **(A)** shows an enriched pathway in which genes that are localised to ribosomes, and the cell envelope are highly expressed. **B)** Proteins that give rise to graph **A**, have been used as input for **B**. Nodes in red represent the proteins used as input for the STRING analysis. The grey nodes represent proteins that interact with the input proteins. Edges represent the interaction and intensity of the edge colour, grey to dark represents the confidence of the interaction.

As shown in FIG 6B, the proteins in red nodes represent 30S ribosomal proteins such as S12/rpsL, S19/rpsS, S20/rpsT and S18/rpsR1, whereas the red nodes-L22/rplV, along with white node-L25/Ctc (rplY) belong to 50S ribosome. Protein for 30S ribosome biogenesis, GTPase-EngC-AKE00588.1, elongation factor-LepA (white node), and assembly factor-RbfA, all three of them act as a quality control for 30S assembly.

On the same line, TatB, a core component of the twin-arginine translocation (Tat) system, participates in exporting folded proteins across the cytoplasmic membrane into the periplasm or cell envelope (65,66). Around this ribosomal core-cell envelope axis, other localized proteins do belong to other biological processes, for instance, i) Proteins involved in nitrogen synthesis (such as N-acyl-L-amino acid amidohydrolase-YxeP_5-(AKD97065.1), succinyldiaminopimelate aminotransferase-DapC-AKD99089.1 and the ACT-domain amino acid-binding sensor-AKD97276.1, ii) Oxido-reductase stress such as the heme peroxidase-AKD98601.1, the ferritin-like diiron hydroxylase-AKE00593.1 and the short-chain dehydrogenase-AKD98624.1, iii) Regulators which sense amino acids and RNA including the UPF0109-family RNA-binding protein-AKD97324.1, the Lrp/AsnC-family transcription factor-AKD98171.1 and the LuxR/NarL-family response regulator-AKD96967.1 and iv) metal homeostasis and catabolic pathway such as the Fur-family metal-responsive regulator-AKE00779.1 and the ferritin-like diiron hydroxylase-AKE00593.1. The Markov (MCL) analysis reinforces the above idea by keeping 27 proteins which are localised to 30S and 50 S ribosomes (see supplementary file X16).

### Dual stress leads to rewiring of proteome function towards efficient translation, environmental sensing and nitrogen scavenging

There is differential enrichment of molecular functions in ShinyGO analysis (FIG 7A and 7 C, for details, see (Supplementary Files X17 and X18). There is enrichment of genes which are involved in ribosome biogenesis and it is represented in FIG 7A as, ‘rRNA binding’, or ‘Structural constituent of ribosome’. Overall, these proteins are involved in ribosomal binding, structure, and translational fidelity. Their STRING analysis give rise to proteins which ultimately protect DNA such as the RecG-like DNA-binding protein-AKD97587.1 and related factor, RNA/ribosomes/transcriptional factors such as the KhpA-like RNA-binding protein-AKD97324.1 and transcriptional regulators-WhiB3/AKD96955.1, the LuxR/NarL-type regulator-AKD96967.1, AsnC/Lrp-family regulator-AKD98171.1, protein foldings including the histidine kinase-KinE-AKD98528.1, associated redox-sensing and metabolic regulators. MCL Clustering (Supplementary File X19) indicates two major changes in gene expression. One large module of proteins includes many 30S/50S proteins and IF-3, as well as one RNA-binding protein (AKD97324.1), together, these help maintain rRNA stability along with high-fidelity translation. A smaller, two-component pair (KinE + LuxR/NarL-type regulator) operates as a middleman between environmental conditions and gene expression. It ensures that only the energy required for DNA repair, transcription, and translation is used in dual-stress conditions.

**FIG 7.**
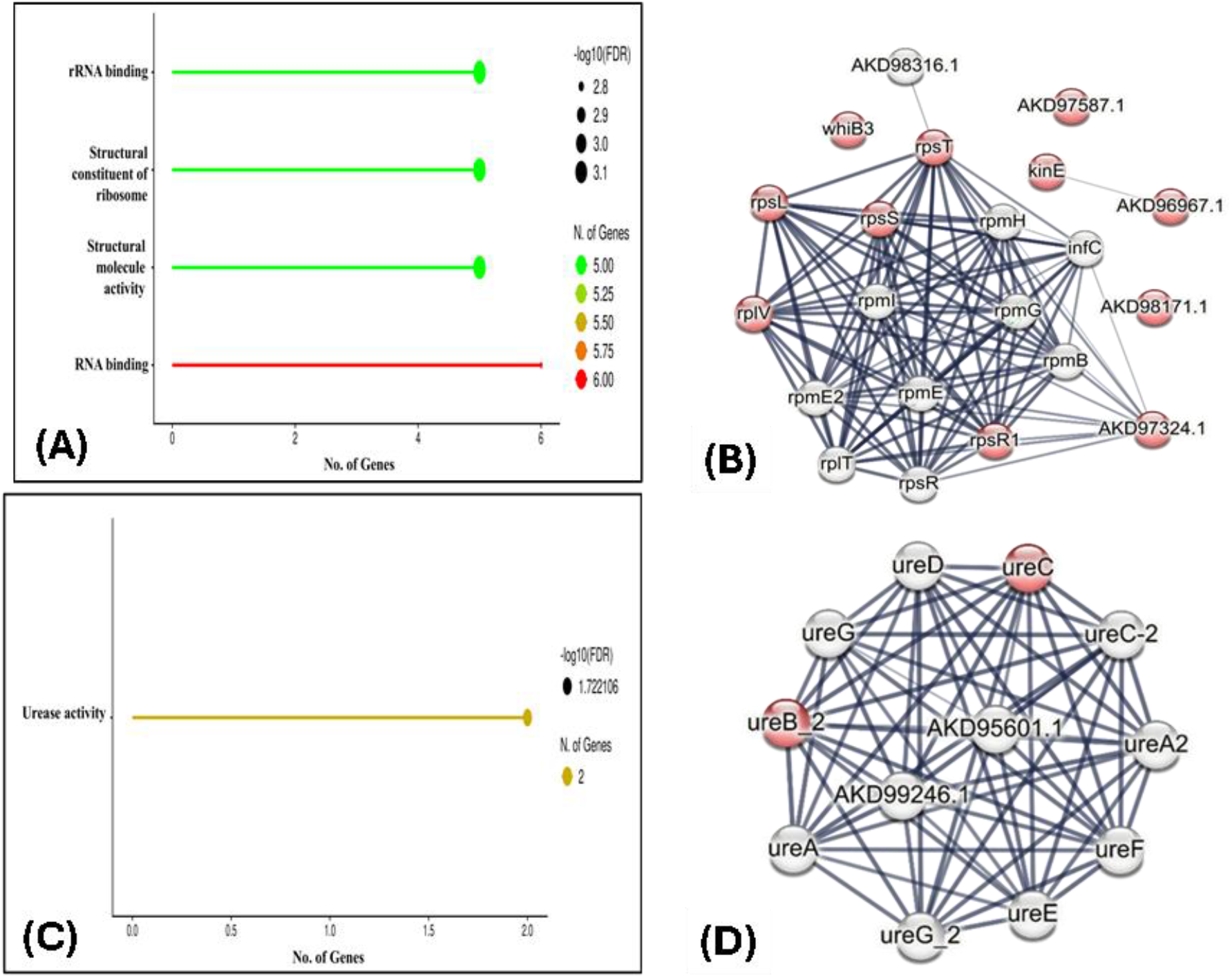
Molecular function enrichment analysis of differentially expressed cytosolic proteins and their PPI (Protein-protein interaction) networks. The legends are like those in Fig. 5, with emphasis on molecular functions. ShinyGO 0.85 is used to analyse molecular functions (58) as shown in A and C. STRING version 12.0 is used for protein-protein interaction (59) as shown in B and D. (**A and C)** Here, the input comes from the upregulated and downregulated data which was obtain from LC-MS/MS analysis. The X axis indicates the number of genes for each ShinyGO molecular function, and the Y axis shows the name of the ShinyGO molecular functions. The graph in (A) shows an enriched set of molecular functions, with genes involved in ribosomes highly expressed. In (C), repressed proteins of dual stress have been used as input and a single enriched molecular function, ‘Urease activity’, is shown on the Y-axis. (**B and D)** proteins that give rise to graph **A**, have been used as input for **B** and proteins that appear in graph **C** have been used as input for **D**. Nodes in red represent the proteins used as input for the STRING analysis. The grey nodes represent proteins that interact with the input proteins. Edges represent the interaction and intensity of the edge colour, grey to dark represents the confidence of the interaction.

There is repression of the whole urease/urea-hydolyzing module. Both catalytic urease subunits (UreC, UreB and the small urease β-like AKD99246.1) plus γ subunits (UreA/A2, UreC-2) and all accessory maturation proteins are equally downregulated. The associated high-affinity urea permease-AKD95601.1 is also part of this module, creating a direct link between import of urea and nitrogen-limiting condition. This string MCL clustering (Supplementary File X20) reinforces the same idea with an emphasis on nickel-centre maturation. The urease enzyme requires Ni^+2^ as a cofactor; if the Ni^+2^ centre is not mature, the enzyme will ultimately not scavenge nitrogen from urea.

## 4. DISCUSSION AND FUTURE PERSPECTIVES

*Rhodococcus sp*. has been used as a bioremediation agent for a long time. In this study, authors are looking for its biofertilizer and PGP in vitro using proteomics analysis under nitrogen-deficient and high-salt conditions. During dual stress, the main adaptations that *Rhodococcus sp* has undergone are described ahead.

### Translational-fidelity is essential under the dual stress

In dual stress, RS1 not only fails to down-regulate growth but also actively remodels its cytoplasmic proteome as reported before in Patrauchan et al., 2012 (12), with about one-quarter of cytoplasmic proteins showing marked changes in abundance. The strong upregulation of ribosomal proteins (RpsL, RpsS, RpsT, RpsR1, RplV, RplL and RplA), elongation factor Tuf and chaperones suggests bacteria are investing in a smaller yet higher-quality translational apparatus (12). This maintains accurate protein synthesis in the face of salt and nitrogen stress. The strategy of optimising performance rather than chasing numbers is bacterial adaptive strategy to stressful environments. Dual stress response prioritise the accurate synthesis of key stress-response and envelope proteins rather than large-scale biomass accumulation.

### Environmental sensors are active in Stress conditions

There are three levels of understanding the adaptation of *Rhodococcus sp*. to nitrogen-deficient and salt-stress conditions. The first one is central metabolism, which represents the normal reactions running in the bacteria, the second is gene expression, which will change as per the survival strategy of the particular bacterium due to its evolutionary history, and the third is environmental status, changing environment will change the stimuli (osmolarity, pH, nitrogen abundance, redox status) to which bacteria will respond. To sense the environment, many proteins have evolved, such as KinE and thiol-linked transcriptional regulators/enzymes such as WhiB3, Hsp18, AhpC, and MetE. WhiB-type regulators are known redox sensors in actinobacteria (67) and their joint upregulation with peroxidases (AhpC) and small heat-shock proteins (Hsp18) indicates that bacteria experience oxidative and proteotoxic stress in our dual stress conditions. An additional hint is, high expression of Dps-DNA protecting protein, which not only binds iron but also protect the genomic DNA, thereby hinting towards the DNA damage due to dual stress.

### Bacterial adaptive strategies: Osmolyte production, PGP-linked proteins and Cell envelope remodelling

RS1 seems to regulate osmotic pressure and cell envelope boundary equilibrium by up-regulating enzymes involved in the synthesis of protective osmolytes such as ectoine (e.g. EctC), endopeptidases (NlpC/P60) and many other lipid/envelope-modifying molecules (e.g. threonine dehydratase IlvA, acetyl-CoA C-acetyltransferase, 2-oxoisovalerate dehydrogenase, cutinases/esterases and lipases). Ectoine is a well-established compatible solute that protects proteins and membranes (68), while NlpC/P60 and lipid-modifying enzymes can manipulate peptidoglycan structure to maintain envelope integrity under high-salt conditions (69). Such traits have excellent potential for PGP biofertilizers in saline agriculture, as they improve rhizosphere persistence and stress resistance and indirectly support plant development. Along this line, STRING and Markov analysis also highlights an axis of ribosome–cell envelope which includes export component like Tat and envelope-associated factors like Soj_3 and hydrolases like PldB.

### Forbidding urease activity channels energy into more efficient pathways

In dual stress, down-regulation of urease structural components (UreA, UreB, UreC, UreE, UreG) and the associated nickel-insertion machinery, as well as high-affinity ammonium-transport systems (GlnK, Amt), suggests that RS1 preferred to shift its energy away from the expensive pathways of nitrogen accumulation (70) to low-energy routes for nitrogen assimilation, such as IlvA-mediated deamination or NAD-dependent glutamate dehydrogenase (71). This results in a direct relevance for the biofertilizer or crop transgenic engineering. The up-regulated cross-tolerance proteins and PGP-linked enzymes such as WhiB3, ectoine synthase, Dps, IlvA and redox enzymes could serve as potential resources for overexpression in plants or beneficial crops, while the genes encoding urease and high-affinity ammonium-transporters seem plausible candidates to be attenuated or knocked out.

In summary, the altered proteomic profile of *Rhodococcus jialingiae* RS1 under dual stress conditions supports its potential as a PGPB and biofertilizer. RS1 fixes the translation and envelope, strictly controls redox and nitrogen economies, produces osmo-protectants under dry conditions and diverts metabolic currency from nitrogen scavenging pathways. These characteristics make it suitable for use as a biofertilizer for soil deficient in nitrogen and enriched in NaCl.

## Acknowledgements

We thank Mr Yash Pratap Rastogi from Central Instrumentation Facility (CIF), University of Delhi South Campus, for guiding us on preparing samples for Orbitrap-derived LC-MS/MS proteomics. PhD committee members of the Department of Bio-Sciences and Technology, MMDU, are highly appreciated for their valuable inputs. The authors are highly thankful to CRL (Central Research Laboratory) and the Director of the Research & Development (R&D) Cell of MMDU (Maharishi Markandeshwar Deemed to be University) for providing the state-of-the-art facilities to carry out this research work. The graphical abstract was created with BioRender (https://BioRender.com/jc0zbwz).

## Author Contribution

AP and BS have directed the research project. Experiments are primarily conducted by SAM, with small contributions from SA, VKM, GKW and AS. AP, SAM, and AKS have written the manuscript, taking input from VKM, SA, GKW and AS. The authors thank Dr Suchitra Upreti, a BIOCARE fellow at ICAR-IVRI, Izzatnagar, for her guidance on improving the manuscript’s flow and grammar.

## Declaration

Authors are not declaring any competing interests.

## Data-Availability

We have deposited the relevant proteomics data files in the open-access communal repository, Mendeley Data. The DOI of the dataset is “10.17632/p3m5v2kxmt.1”. The research conducted in this manuscript is supported by the data included in the main article/supplementary files/Mendeley Database/ files that would be made available on request.

## References

1. Sanower Hossain M, Sultan Ahmad Shah J. Present Scenario of Global Salt Affected Soils, its Management and Importance of Salinity Research ARTICLE INFORMATION [Internet]. Vol. 1, International Research Journal of Biological Sciences Perspective. 2019. Available from: www.scirange.com

2. Ayangbenro AS, Babalola OO. Reclamation of arid and semi-arid soils: The role of plant growth-promoting archaea and bacteria. Curr Plant Biol [Internet]. 2021;25:100173. Available from: 10.1016/j.cpb.2020.100173

3. Fanai A, Bohia B, Lalremruati F, Lalhriatpuii N, Lalrokimi Lalmuanpuii, R, et al. Plant growth promoting bacteria (PGPB)induced plant adaptations to stresses: an updated review. PeerJ. 2024;12(8):1–37.

4. Khumairah FH, Setiawati MR, Fitriatin BN, Simarmata T, Alfaraj S, Ansari MJ. Halotolerant Plant Growth-Promoting Rhizobacteria Isolated From Saline Soil Improve Nitrogen Fixation and Alleviate Salt Stress in Rice Plants. 2022;13(June):1–14.

5. Sunita K, Mishra I, Mishra J, Prakash J, Arora NK. Secondary Metabolites From Halotolerant Plant Growth Promoting Rhizobacteria for Ameliorating Salinity Stress in Plants. 2020;11(October):1–12.

6. Goszcz A, Furtak K, Stasiuk R, Wójtowicz J, Musiałowski M, Schiavon M, et al. Bacterial osmoprotectants — a way to survive in saline conditions and potential crop allies. 2025;(May).

7. Kaleh AM, Singh P, Ooi Chua K, Harikrishna JA. Modulation of plant transcription factors and priming of stress tolerance by plant growth-promoting bacteria: a systematic review. Ann Bot. 2025;135(3):387–402.

8. Bhattacharyya PN, Jha DK. Plant growth-promoting rhizobacteria (PGPR): Emergence in agriculture. Vol. 28, World Journal of Microbiology and Biotechnology. 2012. p. 1327–50.

9. Mohammed SA, Aman S, Singh B. Unveiling the positive impacts of the genus Rhodococcus on plant and environmental health. J Exp Biol Agric Sci. 2024 Sep 25;12(4):557–72.

10. Alvarez HM, Silva RA, Cesari AC, Zamit AL, Peressutti SR, Reichelt R, et al. Physiological and morphological responses of the soil bacterium Rhodococcus opacus strain PD630 to water stress. FEMS Microbiol Ecol. 2004 Nov 1;50(2):75–86.

11. Francis IM, Stes E, Zhang Y, Rangel D, Audenaert K, Vereecke D. Mining the genome of Rhodococcus fascians, a plant growth-promoting bacterium gone astray. N Biotechnol. 2016;33(5):706–17.

12. Patrauchan MA, Miyazawa D, Leblanc JC, Aiga C, Florizone C, Dosanjh M, et al. Proteomic Analysis of Survival of Rhodococcus jostii RHA1 during Carbon Starvation. 2012;78(18):6714–25.

13. Kuhl T, Chowdhury SP, Uhl J, Rothballer M. Genome-Based Characterization of Plant-Associated Rhodococcus qingshengii RL1 Reveals Stress Tolerance and Plant– Microbe Interaction Traits. Front Microbiol [Internet]. 2021 Aug 18;12. Available from: https://www.frontiersin.org/articles/10.3389/fmicb.2021.708605/full

14. Saeki K, Kumagai H. The rnf gene products in Rhodobacter capsulatus play an essential role in nitrogen fixation during anaerobic DMSO-dependent growth in the dark. 1998;464–7.

15. Kosová K, Vítámvás P, Urban MO, Prášil IT, Renaut J. Plant abiotic stress proteomics: The major factors determining alterations in cellular proteome. Front Plant Sci. 2018;9(February):1–22.

16. Jia YL, Chen H, Zhang C, Gao LJ, Wang XC, Qiu L Le, et al. Proteomic analysis of halotolerant proteins under high and low salt stress in Dunaliella salina using two-dimensional differential in-gel electrophoresis. Genet Mol Biol. 2016 Apr 1;39(2):239–47.

17. Panda P, Giri SJ, Sherman LA, Kihara D, Aryal UK. Proteomic changes orchestrate metabolic acclimation of a unicellular diazotrophic cyanobacterium during light-dark cycle and nitrogen fixation states [Internet]. 2024. Available from: http://biorxiv.org/lookup/doi/10.1101/2024.07.30.605809

18. Liu GH, Yang S, Han S, Xie CJ, Liu X, Rensing C, et al. Nitrogen fixation and transcriptome of a new diazotrophic Geomonas from paddy soils. MBio. 2023 Dec 1;14(6).

19. Sharma S, Kulkarni J, Jha B. Halotolerant rhizobacteria promote growth and enhance salinity tolerance in peanut. Front Microbiol. 2016 Oct 13;7(OCT).

20. Wilson ST, Tozzi S, Foster RA, Ilikchyan I, Kolber ZS, Zehr JP, et al. Hydrogen cycling by the unicellular marine diazotroph crocosphaera watsonii strain WH8501∇. Appl Environ Microbiol. 2010 Oct;76(20):6797–803.

21. Yasmin H, Naeem S, Bakhtawar M, Jabeen Z, Nosheen A, Naz R, et al. Halotolerant rhizobacteria Pseudomonas pseudoalcaligenes and Bacillus subtilis mediate systemic tolerance in hydroponically grown soybean (Glycine max L.) against salinity stress. PLoS One [Internet]. 2020;15(4):1–26. Available from: 10.1371/journal.pone.0231348

22. Khumairah FH, Setiawati MR, Fitriatin BN, Simarmata T, Alfaraj S, Ansari MJ, et al. Halotolerant Plant Growth-Promoting Rhizobacteria Isolated From Saline Soil Improve Nitrogen Fixation and Alleviate Salt Stress in Rice Plants. Front Microbiol. 2022 Jun 6;13.

23. Wang Z, Xu J, Li Y, Wang K, Wang Y, Hong Q, et al. Rhodococcus jialingiae sp. nov., an actinobacterium isolated from sludge of a carbendazim wastewater treatment facility. Int J Syst Evol Microbiol. 2010;60(2):378–81.

24. Schober, Patrick MD, PhD, MMedStat; Boer, Christa PhD, MSc; Schwarte, Lothar A. MD, PhD M. Anesthesia & Analgesia. [cited 2026 Apr 21]. p. 1763–8 Correlation Coefficients: Appropriate Use and Interpretation. Available from: https://journals.lww.com/anesthesia-analgesia/fulltext/2018/05000/correlation_coefficientsappropriate_use_and.50.aspx

25. Oliveros JC. Venny 2.1.0: An interactive tool for comparing lists with Venn’s diagrams [Internet]. 2015. Available from: https://bioinfogp.cnb.csic.es/tools/venny/

26. Hu X, Li D, Qiao Y, Song Q, Guan Z, Qiu K, et al. Salt tolerance mechanism of a hydrocarbon-degrading strain : Salt tolerance mediated by accumulated betaine in cells. J Hazard Mater [Internet]. 2020;392(December 2019):122326. Available from: 10.1016/j.jhazmat.2020.122326

27. Schiess R, Mueller LN, Schmidt A, Mueller M, Wollscheid B, Aebersold R. Analysis of cell surface proteome changes via label-free, quantitative mass spectrometry. Mol Cell Proteomics. 2009 Apr;8(4):624–38.

28. Access O. Reproducibility and reliability assays of the gene expression-measurements. 2014;1–10.

29. Cheng-guang H, Gualerzi CO. The Ribosome as a Switchboard for Bacterial Stress Response. 2021;11(January):1–15.

30. Kim T hyung, Leslie P, Zhang Y. Ribosomal proteins as unrevealed caretakers for cellular stress and genomic instability ABSTRACT : 2014;5(4).

31. Bu Y xin, Xu L. Understanding the mechanisms of halotolerance in members of Pontixanthobacter and Allopontixanthobacter by comparative genome analysis. 2023;(March).

32. Pagano E, Ardila F, Ayub D, Mozzicafreddo M, Cuccioloni M, Angeletti M. Acetoacetyl-CoA thiolase regulates the mevalonate pathway during abiotic stress adaptation. 2011;62(15):5699–711.

33. Vejzovic D, Schwaiger T, Topciu A, Petrowitsch L, Arnautovic A. Bacterial cell fate under stress : lipid remodeling and antimicrobial peptide attack. 2026;

34. Suvarna K, Biswas D, Pai MGJ, Acharjee A, Singh A, Banerjee A, et al. Proteomics and Machine Learning Approaches Reveal a Set of Prognostic Markers for COVID-19 Severity With Drug Repurposing Potential. 2021;12(April):1–18.

35. © 1970 Nature Publishing Group. 1970;

36. National Human Genome Research Institute, National Institutes of Health [Internet]. [cited 2026 Apr 18]. Central dogma. Available from: https://www.genome.gov/genetics-glossary/Central-Dogma

37. Starosta AL, Lassak J, Jung K, Wilson DN. The bacterial translation stress response. FEMS Microbiol Rev. 2014;38(6):1172–201.

38. Shahriari F. Effect of salt stress on genes encoding translation-associated proteins in Arabidopsis thaliana. 2012;(September):1095–102.

39. Urwin L, Savva O, Corrigan RM. Microbial Primer : what is the stringent response and how does it allow bacteria to survive stress ? 2024;

40. Nandi SK, Panda AK, Chakraborty A, Rathee S, Roy I. Role of ATP-Small Heat Shock Protein Interaction in Human Diseases. 2022;9(February):1–10.

41. Nandi SK, Chakraborty A, Panda AK, Ray SS. Interaction of ATP with a Small Heat Shock Protein from Mycobacterium leprae : Effect on Its Structure and Function. 2015;1–28.

42. Liu R, Wang S, Song J. New progress in the production, oxidative damage, and scavenging mechanisms of reactive oxygen species in plants under abiotic stress. 2026;(March):1–18.

43. Section IV Crosstalk Among Reactive Oxygen, Nitrogen and Sulfur Species Reactive Oxygen Species, Reactive Nitrogen Species and Oxidative Metabolism Under Waterlogging Stress.

44. Wasim M, Bible AN, Xie Z, Alexandre G, Alexandre G. Alkyl hydroperoxide reductase has a role in oxidative stress resistance and in modulating changes in cell-surface properties in Azospirillum brasilense Sp245. 2009;1192–202.

45. Poole LB. Bacterial defenses against oxidants: mechanistic features of cysteine-based peroxidases and their flavoprotein reductases. 2005;433:240–54.

46. Stadtman ER, Levine RL. Free radical-mediated oxidation of free amino acids and amino acid residues in proteins Opening Review Article. 2003;207–18.

47. Hoshi T, Heinemann SH, Faculty M, Schiller F, Strae D, Jena D. Topical Review Regulation of cell function by methionine oxidation and reduction. 2001;1–11.

48. Kuhlmann AU, Hoffmann T, Bursy J, Jebbar M, Bremer E. Ectoine and Hydroxyectoine as Protectants against Osmotic and Cold Stress : Uptake through the SigB-Controlled Betaine-Choline-Carnitine Transporter-Type Carrier EctT from Virgibacillus pantothenticus M. 2011;193(18):4699–708.

49. Roberts MF. microorganisms. 2005;30:1–30.

50. Guillouet S, Rodal AA, An G, Lessard PA. Expression of the Escherichia coli Catabolic Threonine Dehydratase in Corynebacterium glutamicum and Its Effect on Isoleucine Production. 1999;65(7):3100–7.

51. Geiman DE, Raghunand TR, Agarwal N, Bishai WR. Differential Gene Expression in Response to Exposure to Antimycobacterial Agents and Other Stress Conditions among Seven Mycobacterium tuberculosis whiB-Like Genes. 2006;50(8):2836–41.

52. Ramón-garcía S, Ng C, Jensen PR, Dosanjh M, Burian J, Morris RP, et al. WhiB7, an Fe-S-dependent Transcription Factor That Activates Species-specific Repertoires of Drug Resistance Determinants in Actinobacteria *. 2013;288(48):34514–28.

53. Chawla M, Mishra S, Anand K, Parikh P, Mehta M, Vij M, et al. Redox Biology Redox-dependent condensation of the mycobacterial nucleoid by WhiB4. Redox Biol [Internet]. 2018;19(August):116–33. Available from: 10.1016/j.redox.2018.08.006

54. (PDF) Present Scenario of Global Salt Affected Soils, its Management and Importance of Salinity Research [Internet]. [cited 2026 Feb 14]. Available from: https://www.researchgate.net/publication/334773002_Present_Scenario_of_Global_Salt_Affected_Soils_its_Management_and_Importance_of_Salinity_Research

55. Proteau PJ. reductoisomerase : an overview. 2004;32:483–93.

56. Chem JB. Papers of the Week Crystal Structure of Bacterial Enzyme Reveals a Potential Target for New Antibiotics 354811;2012.

57. Education N. Scitable by Nature Education. 2014 [cited 2026 Apr 18]. Ribosomes, transcription, and translation. Available from: https://www.nature.com/scitable/topicpage/ribosomes-transcription-and-translation-14120660/

58. Ge SX. ShinyGO : a graphical gene-set enrichment tool for animals and plants. 2020;36(December 2019):2628–9.

59. Szklarczyk D, Kirsch R, Koutrouli M, Nastou K, Mehryary F, Hachilif R, et al. The STRING database in 2023 : protein – protein association networks and functional enrichment analyses for any sequenced genome of interest. 2023;51(November 2022):638–46.

60. Szklarczyk D, Gable AL, Lyon D, Junge A, Wyder S, Huerta-cepas J, et al. STRING v11 : protein – protein association networks with increased coverage, supporting functional discovery in genome-wide experimental datasets. 2019;47(November 2018):607–13.

61. Mols M, Abee T. Role of Ureolytic Activity in Bacillus cereus Nitrogen Metabolism and Acid Survival M. 2008;74(8):2370–8.

62. Konieczna I, Kwinkowski M, Kolesi B, Kami Z, Kaca W. Bacterial Urease and its Role in Long-Lasting Human Diseases. 2012;789–806.

63. Carvalho CCCR De, Fischer MA, Kirsten S, Würz B, Wick LY, Heipieper HJ. Adaptive response of Rhodococcus opacus PWD4 to salt and phenolic stress on the level of mycolic acids. AMB Express. 2016;

64. Paul D. Review Osmotic stress adaptations in rhizobacteria. 2012;1–10.

65. Rose P, Mu M, Fro J. Twin-arginine-dependent translocation of folded proteins. 2012;1029–46.

66. Buck E De, Lammertyn E, Anne J. The importance of the twin-arginine translocation pathway for bacterial virulence. 2008;(August).

67. Griffin ME, Klupt S, Espinosa J, Hang HC. host interactions. 2024;

68. Kuhlmann AU, Bremer E. Osmotically regulated synthesis of the compatible solute ectoine in Bacillus pasteurii and related Bacillus spp. Appl Environ Microbiol. 2002;68(2):772–83.

69. Griffin ME, Klupt S, Espinosa J, Hang HC. Review Peptidoglycan NlpC / P60 peptidases in bacterial physiology and host interactions. Cell Chem Biol [Internet]. 2022;30(5):436–56. Available from: 10.1016/j.chembiol.2022.11.001

70. Collins CM, Orazio SEFD. MicroReview Bacterial ureases : structure, regulation of expression and role in pathogenesis. 1993;9:907–13.

71. Heeswijk WC Van, Westerhoff H V, Boogerd C. Nitrogen Assimilation in Escherichia coli : Putting Molecular Data into. 2013;51.

